# TGF-β downregulates antigen processing and presentation genes and MHC I surface expression through a Smad3-dependent mechanism

**DOI:** 10.1101/2023.01.30.526196

**Authors:** Alix K. Berglund, Anna L. Hinson, Lauren V. Schnabel

## Abstract

Regulation of MHC I surface expression by TGF-β is important for controlling cell-mediated immune responses, but how TGF-β downregulates MHC I is unknown. We investigated this mechanism using flow cytometry and RNA-sequencing to identify the major TGF-β signaling pathway and target genes involved. All three isoforms of TGF-β were found to have similar abilities to downregulate constitutive MHC I surface expression. Inhibiting the type I TGF-β receptor, ALK5, as well the canonical TGF-β signaling molecule Smad3, prevented TGF-β from downregulating MHC I surface expression. RNA-sequencing from horse and human mesenchymal stem cells revealed that multiple genes associated with antigen processing and presentation were downregulated in TGF-β-treated cells with B2M and ERAP1 being downregulated in both species. While downregulation of MHC I surface expression a slow process that continued past 24 hours after TGF-β treatment, downregulation of gene expression of ERAP1 occurred within 12 hours after treatment. B2M expression was not consistently downregulated until after 24 hours and TAP2 expression was inconsistent over a 72-hour period, although it was significantly downregulated at 12 hours after TGF-β treatment. This data supports that TGF-β downregulates antigen processing and presentation genes resulting in decreased surface expression of MHC I through a Smad3-dependent mechanism.

## 1 Introduction

Major histocompatibility complex (MHC) class I molecules are cell surface glycoproteins that initiate cell-mediated immune responses by presenting intracellular peptides to CD8^+^ T cells. MHC I molecules plan an important role in the immune detection and cytotoxicity of cells infected with viruses, intracellular bacteria, neoplastic cells, and allogeneic MHC-mismatched cells. Almost all nucleated cells constitutively express MHC I, but expression is tightly regulated to prevent excessive or inappropriate immune responses (van den Elsen, 2011). For stable MHC I molecules to be presented on the cell surface, molecules must be loaded with cytosolic peptides in the endoplasmic reticulum (ER) and non-covalently bind a β2-microglobulin molecule before being shuttled to the cell membrane (Wieczorek et al., 2017). Additional proteins involved in MHC I expression include TAP1 and TAP2, which form the TAP complex to transport peptides from the cytosol into the ER, ERAP1, which cleaves peptides to 8-10 residues prior to peptide loading, and tapasin or TAP binding protein, which associates the MHC I molecule in proximity to the TAP complex. Mutations or dysregulation of β2-microglobulin or other proteins associated with antigen processing and presentation affect the ability of a cell to express stable MHC I molecules on the cell surface (Zijlstra et al., 1990; Salazar-Onfray et al., 1997; Wang et al., 2013).

Cytokines positively and negatively regulate MHC I surface expression. MHC I, β2-microglobulin, and other antigen processing and presentation genes contain an interferon response element in their promoter regions ensuring that antigen presentation increases during infection or inflammatory responses ((Gobin et al., 1997, 2003; van den Elsen et al., 2004). Treatment of cells with exogenous IFN-g results in rapid upregulation of MHC I gene and surface expression (Rosa et al., 1986; Chan et al., 2008). Opposing IFN-g, TGF-β is known to negatively regulate MHC I surface expression. The TGF-β family consists of three isoforms: TGF-β1, TGF-β2, and TGF-β3. TGF-β1 is the primary isoform expressed by immune cells and loss of TGF-β1 in mice results in a severe autoinflammatory disorder in part due to increased expression of MHC I and MHC II on antigen presenting cells (Geiser et al., 1993; Letterio et al., 1996). TGF-β2 is found at high concentrations in immune privileged sites including the anterior chamber of the eye, hair follicle, and brain and is hypothesized to help prevent autoantigen presentation and maintain immune tolerance (Unsicker et al., 1991; Niederkorn, 2003; Vincze et al., 2010). TGF-β1 and TGF-β2 have been reported to downregulate constitutive and IFN-g-induced MHC class I and MHC class II surface expression (Donnet-Hughes et al., 1995; Ma and Niederkorn, 1995; Berglund et al., 2017). The mechanism by which TGF-β regulates MHC I expression has not yet been established, but TGF-β1 and TGF-β2 downregulate MHC class II gene expression by inhibiting its master transcription factor, the class II transactivator (CIITA), through a Smad3-dependent mechanism (Lee et al., 1997; Dong et al., 2001; Romieu-Mourez et al., 2007). Based on in vivo knockout studies, regulation of MHC molecules by TGF-β appears to play an important role in modulating adaptive immune responses. Elucidating this mechanism is therefore important to understanding the role of TGF-β in regulating MHC I and antigen presentation under normal and disease conditions.

In this study, we investigated the mechanism by which TGF-β decreases MHC I surface expression using bone marrow-derived stem cells (MSCs) from horses and humans. Horses, like humans, are an outbred population and display individual variation in MHC I expression levels similar to similar to humans (Berglund et al., 2017). Additionally, large volumes of bone marrow can be easily and repeatedly obtained from horses to isolate primary MSCs, which are of recent scientific interest due to their immunomodulatory role in tissue healing and cancer (Caplan and Sorrell, 2015; Ridge et al., 2017). We report that TGF-β downregulates MHC I surface expression through canonical Smad3 signaling and that B2M and ERAP1, two genes associated with antigen processing and presentation, are also downregulated by TGF-β.

## 2 Materials and Methods

### 2.1 MSC isolation and culture

MSCs were isolated from the bone marrow of 14 Thoroughbred or Thoroughbred cross horses between the ages of 5 and 16. An equal number of mares and geldings were used in each experiment. Bone marrow aspirates were collected aseptically from the sterum of horses using an 11-gauge Jamshidi biopsy needles under standing sedation with local anesthesia or immediately after anesthesia. Bone marrow aspirates were purified via Ficoll-Paque Plus (GE Healthcare) gradient centrifugation. Isolated cells were cultured at 37 °C unless specified in standing media consisting of 1 g/dl glucose DMEM (Corning), 10% fetal bovine serum (FBS)(Cytivia), 2 mM L-glutamine, 100 U/ml penicillin and streptomycin, and 1 ng/ml recombinant human basic fibroblast growth factor (bFGF)(R&D Systems). Human MSCs were obtained from ATCC (catalog #PCS-500-012) from four donors aged 19 to 25. MSCs from two males and two females were used for each experiment. Human MSCs were cultured in the same media as equine MSCs except with 3 ng/ml bFGF. Equine and human MSCs were passaged with Accutase cell-dissociation solution (Innovative Cell Technologies) and plated at a density of 6,250 cells per cm^2^ on tissue culture plates.

MSCs were treated with TGF-β isoforms by adding 1 ng/ml recombinant human TGF-β1, TGF-β2, or TGF-β3 (Biolegend) to the media for a minimum of 72 hours unless otherwise stated. For IFN-g stimulated assays, untreated MSCs were plated in standard media or media containing the appropriate TGF-β isoform 18 hours prior to the addition of IFN-g to the media. Media were exchanged after 48 hours with fresh standard media or TGF-β media with or without IFN-g. TGF-βRI (ALK5) inhibitor SB525334 (10 μM; Selleckchem) or SIS3-HCl (10 μM; Selleckchem) were added into the indicated culture media. For SIS3 experiments, MSCs were pre-incubated with SIS3-HCl in PBS for 1 hour at 37 °C prior to the addition of media. Media were exchanged at 48 hours with fresh media and inhibitor.

### 2.2 Splenocyte isolation and culture

Splenocytes were isolated by pushing the spleens of 6–9-week-old C57BL/6J mice through a 70-micron filter using the plunger of a 3 ml syringe. Splenocytes were cultured for 5 days in RPMI-1640 (Gibco), 10% FBS, 2 mM L-glutamine, 100 U/ml penicillin and streptomycin, 0.1 mM 2-mercaptoethanol, and 25 U/ml human recombinant IL-2 (PeproTech). Splenocytes were cultured with 5 μg/ml Concanavalin A (ConA) (Sigma-Aldrich) and 2 ng/ml human recombinant TGF-β2 for five days.

### 2.3 Flow cytometry

Cells were washed in PBS and incubated with 10% goat serum prior to labeling with primary antibody. Equine MSCs were labeled with mouse anti-horse MHC I or MHC II antibody (clones cz3 and cz11 respectively, gift from Dr. Doug Antczak) followed by APC-conjugated goat anti-mouse IgG antibody (BD Biosciences). Human MSCs were labeled with APC-conjugated mouse anti-human HLA-ABC (BD Biosciences). 4’,6’-diamidino-2-phenylindole (DAPI) was added at 500 ng/ml 15 minutes prior to analysis to identify dead cells. Fluorescence of live cells was measured using a LSRII flow cytometer equipped with FACSDiva analysis software (BD Biosciences). Data was analyzed using FlowJo v.10 (FlowJo, LLC).

### 2.4 RNA-sequencing

Total RNA was extracted from MSCs using the RNeasy Mini kit (Qiagen, Germantown, MD, USA). Libraries were generated and poly(A) enriched using 1 ug of RNA as input. Indexed samples were sequenced using a 150 base pair paired-end protocol via the HiSeq 2500 (Illumina) according to the manufacturer’s protocol. Sequence reads were trimmed to remove possible adapter sequences and nucleotides with poor quality using Trimmomatic v.0.36. The trimmed reads were mapped to the Equus caballus EquCab 3.0 or Homo sapiens GRCh38 reference genome available on ENSEMBL using the STAR aligner v.2.5.2b. Unique gene hit counts were calculated by using featureCounts from the Subread package v.1.5.2. Using DESeq2, a comparison of gene expression between the untreated and TGF-β2-treated MSCs was performed. The Wald test was used to generate *p*-values and log2 fold changes. Genes with an adjusted *p*-value < 0.05 and absolute log2 fold change > 1 were identified as differentially expressed genes for each comparison. The quantification and poly(A) selection of mRNA, library preparation, sequencing, and RNA-seq bioinformatics were outsourced to GENEWIZ, Inc.

### 2.5 Real-time quantitative PCR

Total RNA was extracted from cells using the RNeasy Mini kit. Generation of cDNA was performed using the iScript cDNA synthesis kit (Bio-Rad). Transcripts were quantified by real-time PCR analysis using the PowerUp SYBR Green (Applied Biosystems) and the QuantStudio 6 Flex (Applied Biosciences). mRNA expression in MSCs was normalized to HPRT1 expression and fold change was calculated as the relative gene expression (2^-ΔΔCt^). Primers were generated using Primer-BLAST unless otherwise specified with sequences and references listed in Table 1. All primers were purchased from IDT Technologies. For all reactions, each condition was performed in triplicate.

**Table 1:**
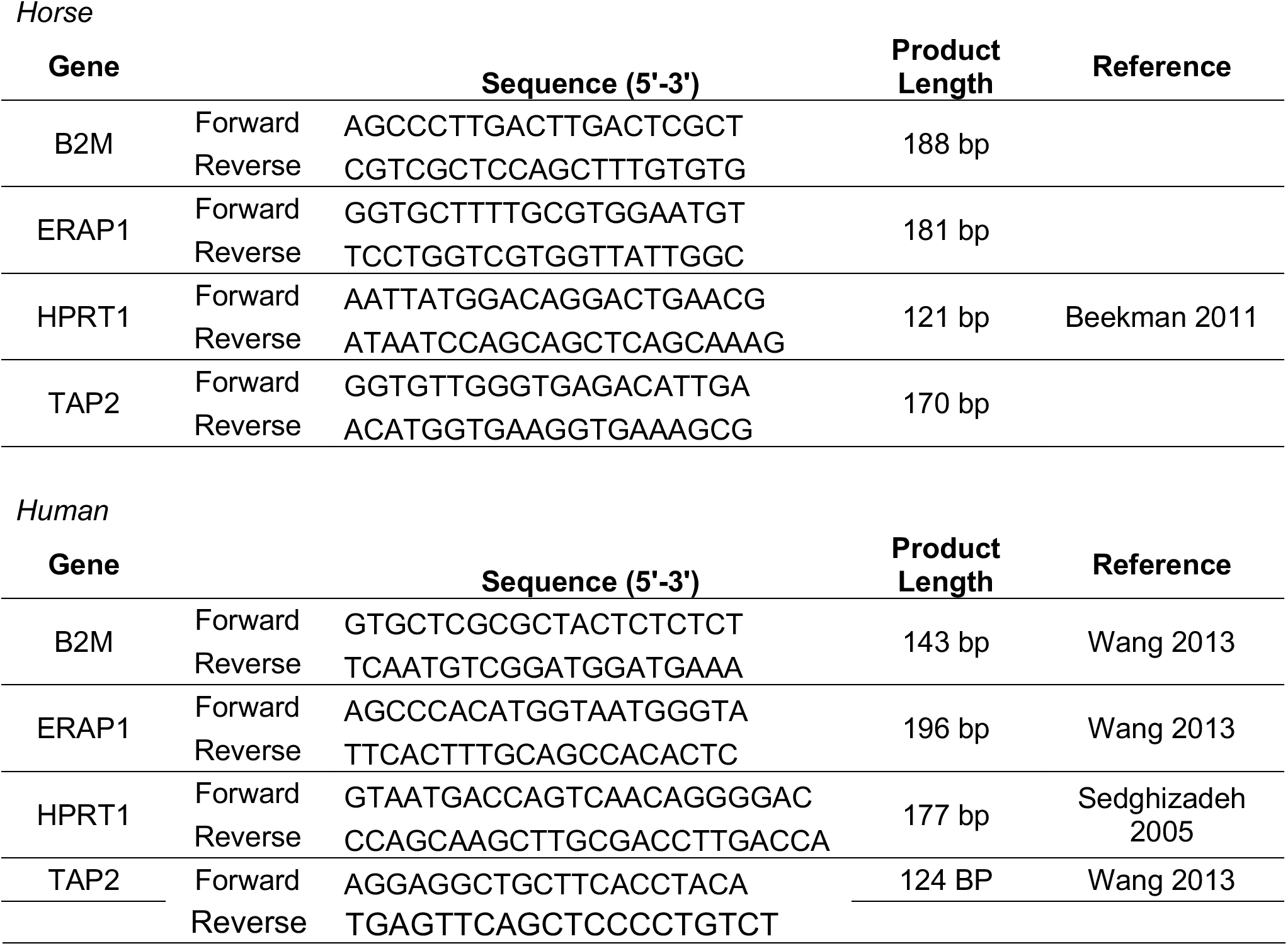
qPCR Primers.

### 2.6 Statistical analysis

Flow cytometry and qPCR data were normalized by log transformation and analyzed by a one or two-tailed paired t-test for or ANOVA using JMP Pro 15 (SAS Institute Inc.). When an ANOVA indicated significant differences (*p* < 0.05), a Tukey’s test was used for multiple comparisons of individual means.

## 3 Results

### 3.1 TGF-β isoforms downregulate constitutive MHC I surface expression

While TGF-β1 and TGF-β2 are known to downregulate MHC I (Ma and Niederkorn, 1995; Berglund et al., 2017), it is unknown if TGF-β3 regulates MHC I expression nor have all three TGF-β isoforms been directly compared in their ability to regulate constitutive and IFN-g-induced MHC I surface expression. All three TGF-β isoforms downregulated constitutive MHC I surface expression and there was no significant difference between the three isoforms (Figure 1a and 1c). None of the three isoforms were effective at blocking IFN-γ-induced MHC I surface expression under these culture conditions as there was no significant difference in the relative GMFI between the untreated and TGF-β-treated groups (Figure 1b and 1c). As all three isoforms were similar in their ability to downregulate constitutive MHC I expression, we continued using TGF-β2 in all subsequent experiments to determine how TGF-β downregulates constitutive MHC I surface expression.

**Figure 1.**
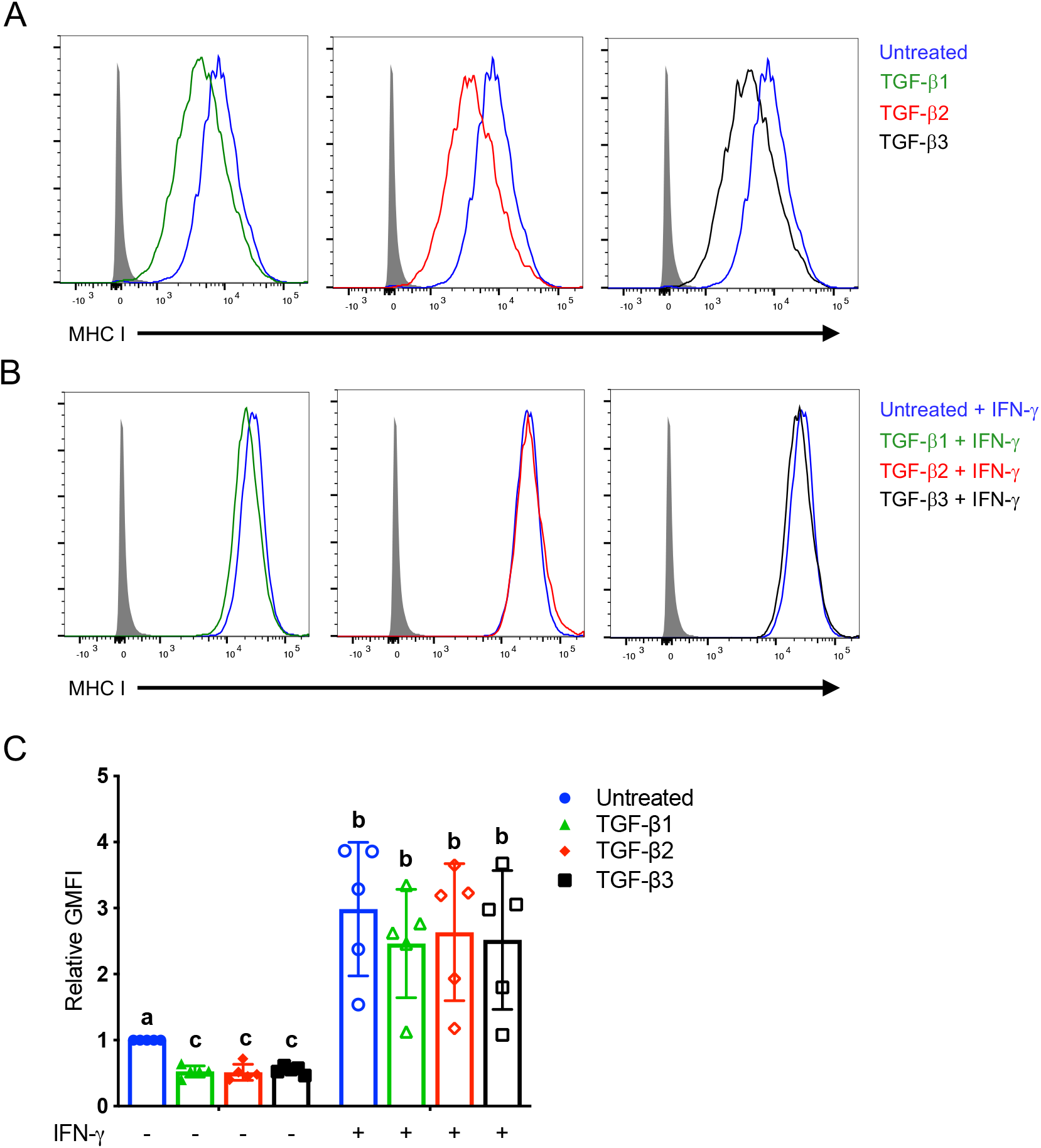
TGF-β isoforms downregulate constitutive MHC I surface expression. **(A)** Representative histograms from one horse depicting MHC I surface expression on MSCs treated with TGF-β1, TGF-β2, or TGF-β3 compared to untreated. **(B)** Representative histograms from one horse depicting IFN-γ-induced MHC I surface expression on MSCs treated with TGF-β1, TGF-β2, or TGF-β3 compared to untreated. **(C)** Relative GMFI of MHC I surface expression on MSCs following TGF-β treatment with or without IFN-γ stimulation. Data shown are mean +/− SD for n = 5 relative to the unstimulated and untreated MSCs. Superscript letters indicate significant differences between groups, *p* < 0.0001 by ANOVA.

### 3.2 TGF-β downregulates MHC I surface expression on human MSCs and mouse splenocytes

We previously published that equine MSCs cultured with TGF-β had significantly decreased MHC I surface expression compared to untreated MSCs (Berglund et al., 2017, 2021). To compare the ability of TGF-β to downregulate MHC I expression across species and cell types, we compared surface expression of MHC I on human MSCs and mouse splenocytes. Human MSCs cultured with TGF-β had a 2.56-fold decrease in MHC I surface expression (Figure 2a). Mouse splenocytes treated with TGF-β also had a 1.98 and 1.82-fold decrease in constitutive H-2K and H-2D surface expression respectively (Figure 2b). Although both H-2K and H-2D expression was upregulated by ConA, splenocytes treated with TGF-β still had a 1.47 and 1.98-fold decrease in H-2K and H-2D surface expression relative to untreated controls (Figure 2c). This supports that regulation of MHC I surface expression by TGF-β is conserved across mammalian species and cell types.

**Figure 2.**
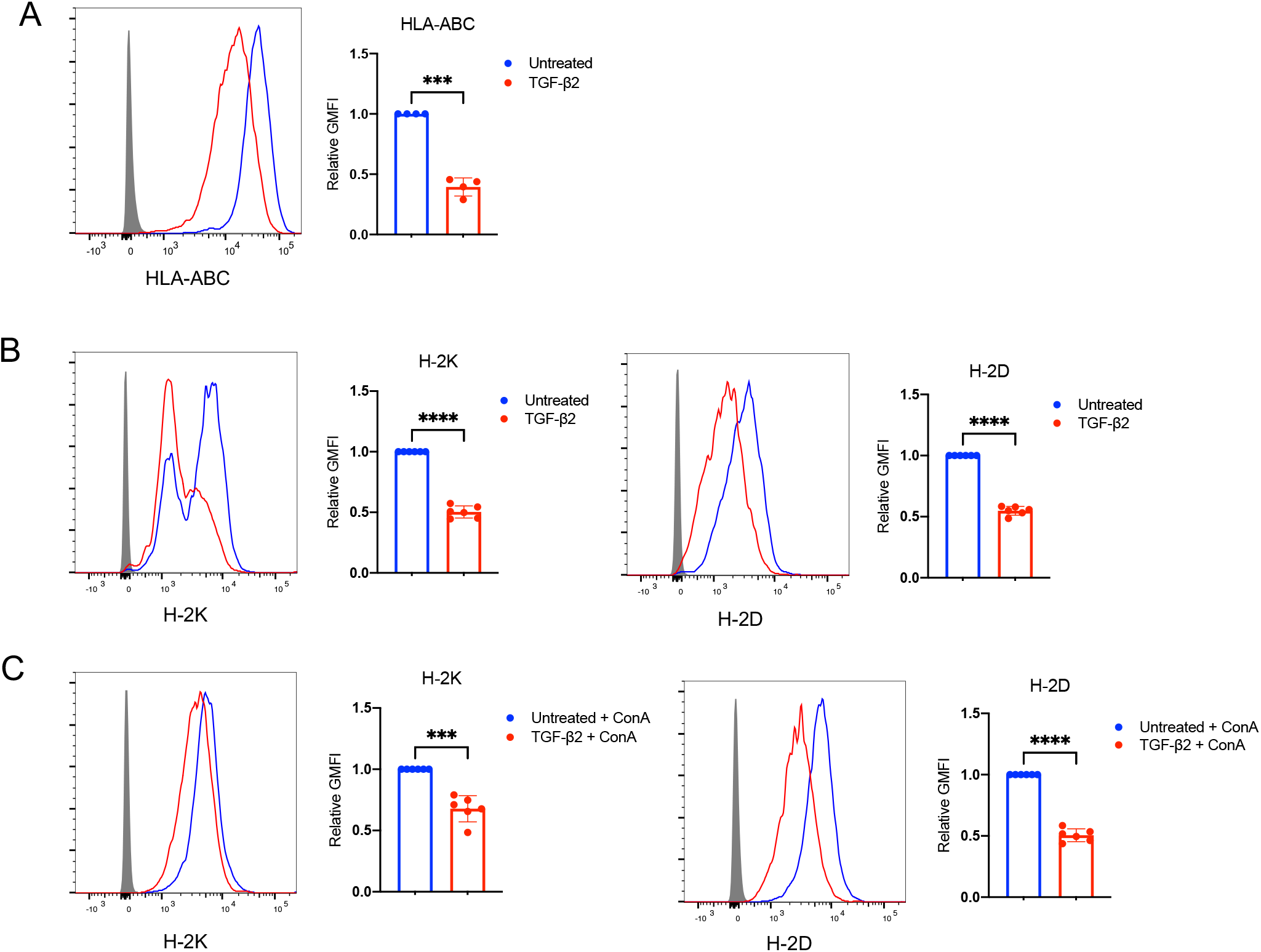
Regulation of MHC I surface expression by TGF-β is conserved across mammals and cell types. **(A)** Representative histogram of HLA-ABC surface expression and relative GMFI of HLA-ABC surface expression on human MSCs treated with TGF-β. Data shown are mean +/− SD for n = 4, *p* = 0.0003 by t-test. (B) Representative histograms of H-2K and H-2D surface expression and relative GMFI of H-2K and H-2D surface expression on mouse MSCs. Data shown are mean +/− SD for n = 6, \*\*\*\**p* < 0.0001 by t-test. (C) Representative histograms for H-2K and H-2D surface expression and relative GMFI of H-2K and H-2D surface expression on mouse splenocytes treated with TGF-β with ConA stimulation. Data shown are mean +/− SD for n = 6, \*\*\**p* = 0.0007, \*\*\*\**p* < 0.0001.

### 3.3 Downregulation of MHC I surface expression by TGF-β is ALK5 and Smad3 dependent

TGF-β isoforms bind to a cell surface TGF-β type II receptor, which then recruits a TGF-β type I receptor. Upon activation, the type I receptor then phosphorylates downstream signaling molecules. ALK5 is one of the two type I receptors and is ubiquitously expressed across cell types (Heldin and Moustakas, 2016). To determine if TGF-β downregulates MHC I surface expression through the ALK5 receptor, MSCs were treated with TGF-β in the presence of the selective ALK5 inhibitor, SB525334. Inhibition of the ALK5 receptor completely abolished the ability of TGF-β to downregulate MHC I surface expression (Figure 3a) confirming that this pathway involves ALK5. Interestingly, inhibition of the ALK5 receptor also increased MHC I surface expression on both untreated and TGF-β-treated MSCs.

**Figure 3.**
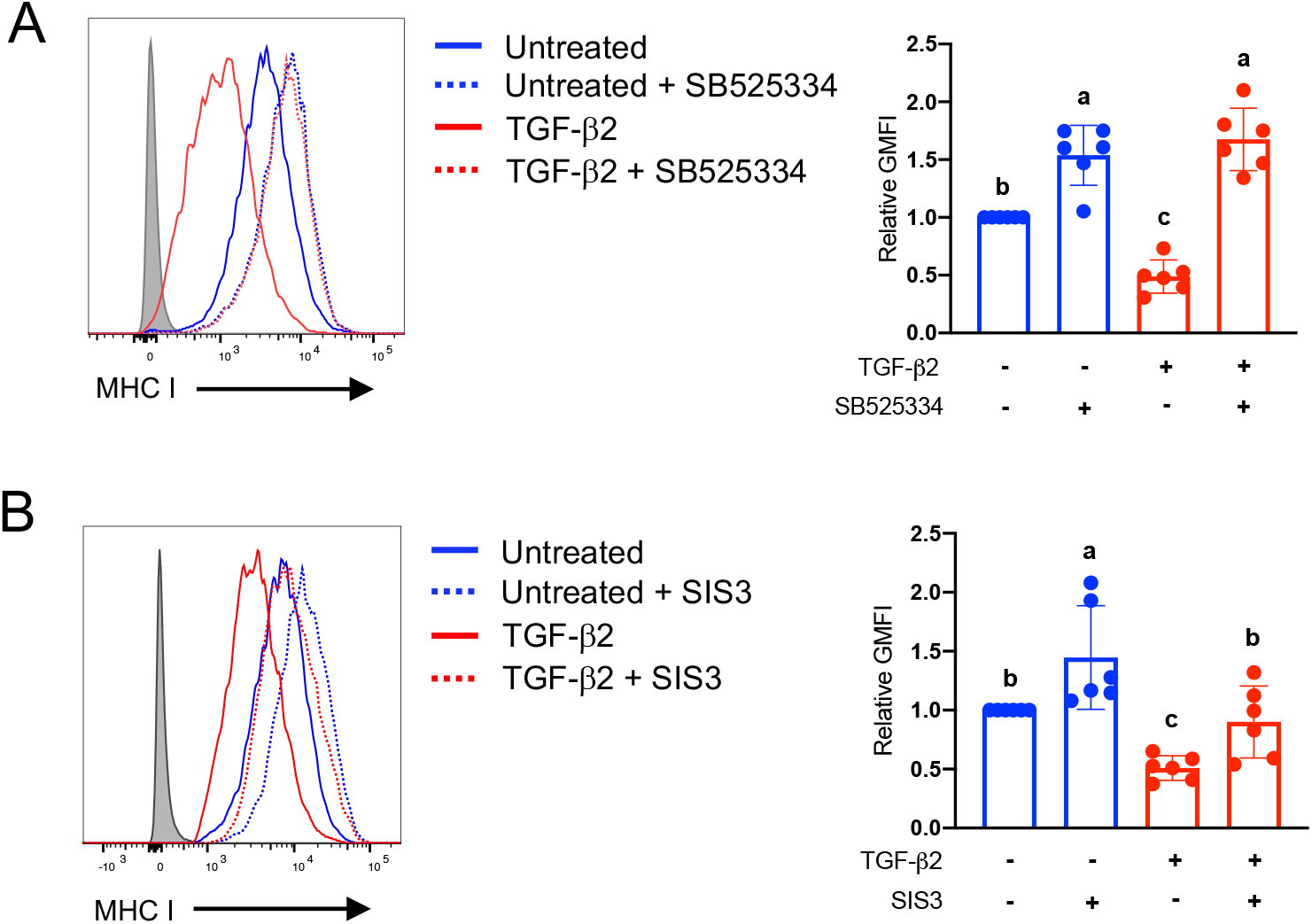
TGF-β downregulates MHC I surface expression through ALK5 and Smad3. **(A)** Representative histogram of MHC I surface expression and relative GMFI of MHC I surface expression on horse MSCs treated with TGF-β with or without an ALK5 inhibitor, SB525334. Data shown are mean +/− SD for n = 6. Superscript letters indicate significant differences between groups, *p* < 0.0001 by ANOVA. **(B)** Representative histogram of MHC I surface expression and relative GMFI of MHC I surface expression on equine MSCs treated with TGF-β with or without a selective Smad3 inhibitor, SIS3. Data shown are mean +/− SD for n = 6. Superscript letters indicate significant differences between groups, *p* < 0.0001 by ANOVA.

Activation of ALK5 results in phosphorylation and activation of Smad2 or Smad3, referred to as the canonical TGF-β signaling pathway, or non-Smad signaling molecules, which are termed non-canonical TGF-β signaling pathways. TGF-β-mediated downregulation of MHC II is known to be through a Smad3-dependent mechanism so we examined if MHC I surface expression is also regulated through Smad3 (Dong et al., 2001). Treatment of MSCs with SIS3-HCl, a selective Smad3 inhibitor, significantly inhibited the ability of TGF-β to downregulate of MHC I surface expression (Figure 3b) supporting that like MHC II, downregulation of MHC I is also via a Smad3-dependent mechanism.

### 3.4 Antigen processing and presentation genes are differentially expressed in untreated and TGF-β-treated MSCs

After phosphorylation of Smad3 by the TGF-β receptor, Smad3 travels to the nucleus and acts as a transcription factor where it positively or negatively regulates transcription depending on which cofactors are bound (Miyazono et al., 2018). To identify target genes of TGF-β for regulating MHC I surface expression, we compared the gene expression of untreated and TGF-β-treated equine and human MSCs using RNA-sequencing. Gene expression of proteins associated with antigen processing and presentation including B2M, ERAP1, TAP2, TAPBP, and TAPBPL were downregulated in TGF-β-treated MSCs, although the degree of downregulation differed between species (Figure 4a and 4b). Expression of MHC I and NLRC5 genes also trended downwards in TGF-β-treated MSCs from both species. Expression of calnexin, calreticulin, ERp57, and proteosome subunits were not differentially expressed in either species (data not shown). qPCR from cell cultures separate from the cultures used for RNA-sequencing confirmed that B2M and ERAP1 gene expression was significantly downregulated in TGF-β-treated MSCs in both species, but a larger sample size demonstrated that TAP2 was downregulated in some, but not all equine and human MSCs (Figure 4c and 4d). Therefore, while there was an overall trend that genes associated with antigen processing and presentation were downregulated, B2M and ERAP1 were the genes most consistently downregulated by TGF-β in both species.

**Figure 4.**
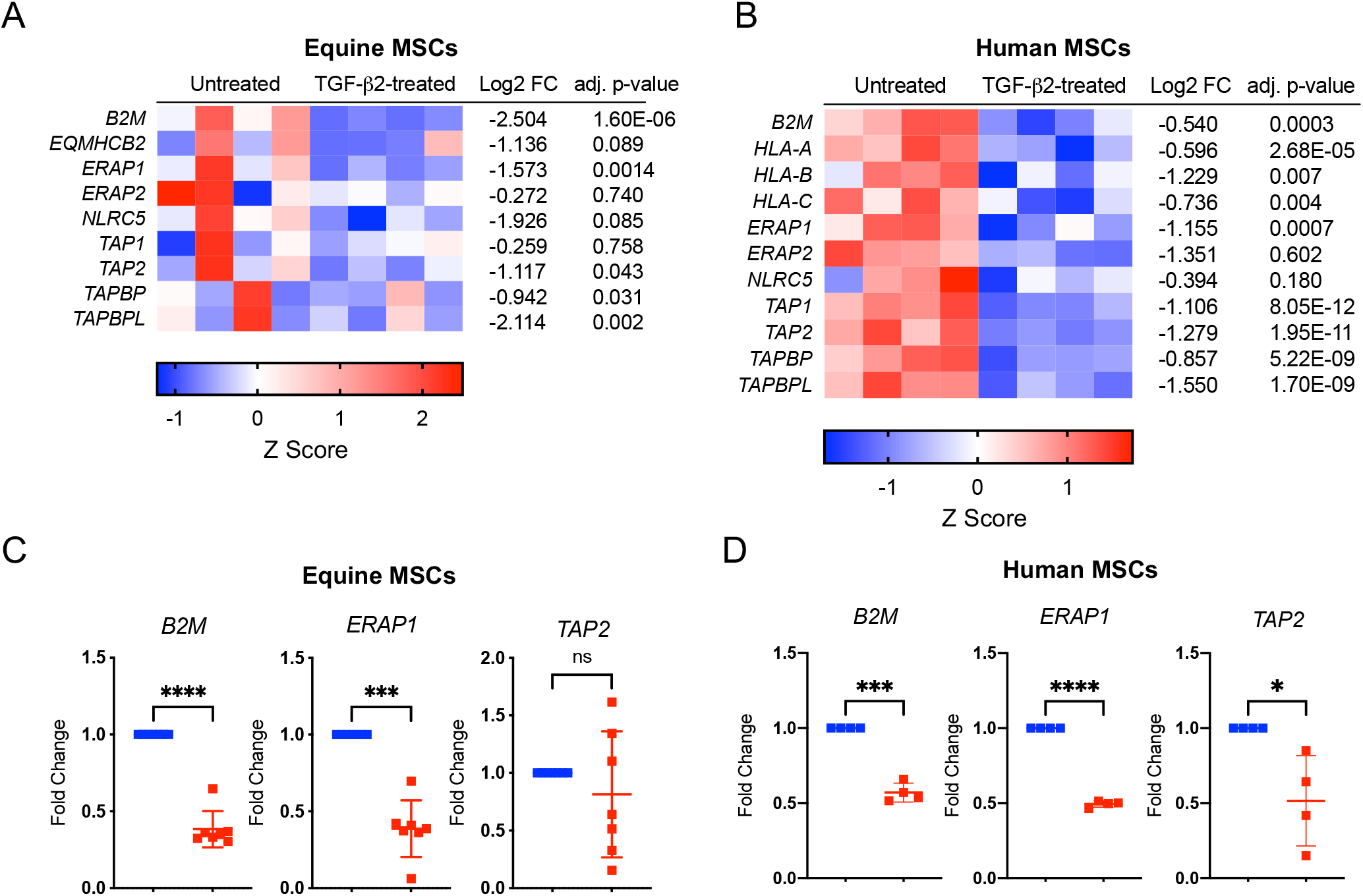
TGF-β downregulates antigen processing and presentation genes. Heatmap representation of antigen processing and presentation genes for untreated and TGF-β-treated **(A)** equine and **(B)** human MSCs. Data shown are z scores, log2 fold change, and the adjusted *p* value for each gene for n = 4. Normalized fold change in gene expression for B2M, ERAP1, and TAP2 in untreated and TGF-β-treated **(C)** equine and **(D)** human MSCs. Data shown are mean +/− SD for n = 7 (horses) and n = 4 (humans). \*\*\*\**p* < 0.0001, \*\*\**p* < 0.001, and \**p* < 0.05 by paired t-test.

### 3.5 Downregulation of MHC I surface expression after TGF-β treatment is progressive

To determine the kinetics of downregulation of MHC I surface expression by TGF-β, MHC I surface expression was measured at 24, 48, and 72 hours after TGF-β treatment. Following treatment with TGF-β, there was a slight, but non-significant decrease in MHC I surface expression by 24 hours, with significant downregulation of MHC I continuing over the next 48 hours (Figure 5a). Gene expression of B2M, ERAP1, and TAP2 were then measured by qPCR up to 72 hours after treatment with TGF-β. B2M expression was significantly lower in TGF-β-treated MSCs at 48- and 72-hours post-treatment while ERAP1 expression was significantly decreased in TGF-β-treated MSCs by 24 hours (Figure 5b). Similar to what was seen in Figure 4, TAP2 expression in TGF-β-treated MSCs was variable between horses, although significant differences were detected at 24 and 72 hours after treatment (Figure 5b). In the ERAP1 and TAP2 data, a noticeable decrease in gene expression was detected in both untreated and TGF-β-treated MSCs at 12 hours, which was determined to be due to the addition of fresh serum to the culture during the media exchange. A separate experiment to measure B2M, ERAP1, and TAP2 expression at 12 hours post-treatment was then conducted where TGF-β was added to the MSC cultures in 100 ul of DMEM without FBS for a final concentration of 1 ng/ml or 100 ul of DMEM alone for the untreated samples. Gene expression of untreated and TGF-β-treated samples were then compared to gene expression in control samples without any additional DMEM added. At 12 hours post-TGF-β-treatment, B2M, ERAP1, and TAP2 expression were significantly decreased compared to untreated samples, although ERAP1 and TAP2 were more consistently downregulated in MSCs compared to B2M (Figure 5c). These results indicate that downregulation of MHC I surface expression by TGF-β is a slow and progressive process that is likely driven by earlier downregulation of antigen processing and presentation genes like B2M and ERAP1.

**Figure 5.**
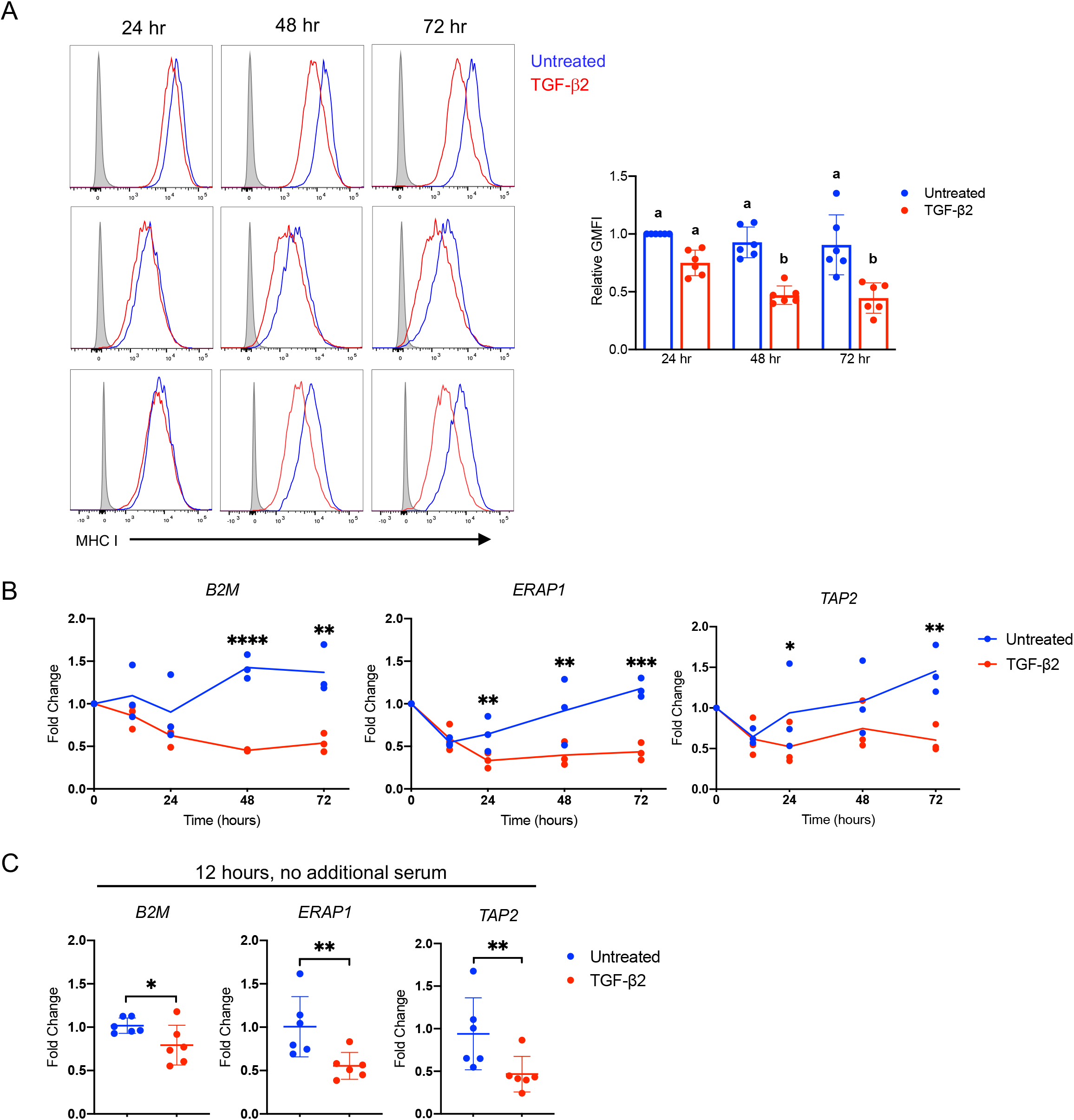
Time course of MHC I surface and antigen processing and presentation gene downregulation by TGF- **(A)** Representative histograms of MHC I surface expression on MSCs from three horses at 24, 48, and 72 hours after TGF-treatment and the relative GMFI at each time point. Data shown are mean +/− SD for n = 6. Superscript letters indicate significant differences between groups, *p* < 0.0001 by ANOVA. **(B)** Normalized fold change for B2M, ERAP1, and TAP2 in equine MSCs at 12, 24, 48, and 72 hours after TGF-treatment. Data shown are mean +/− SD at each time point for n = 3. For B2M, \*\*\*\**p* < 0.0001, \*\**p* = 0.002; ERAP1, \*\**p* < 0.01, \*\*\**p* = 0.0003; TAP2, \**p* = 0.47, \*\**p* = 0.0038 by one-tailed paired t-test. **(C)** Normalized fold change for B2M, ERAP1, and TAP2 in equine MSCs at 12 hours when TGF-β was added without serum. Data shown are mean +/− SD for n = 6. For B2M, \**p* = 0.0147; ERAP1, \*\**p* = 0.0016; TAP2 \*\**p* = 0.0029 by one-tailed paired t-test.

## Discussion

TGF-β is a pivotal regulator of immune and inflammatory responses. The roles of TGF-β in maintaining immune tolerance and immune cell development have been extensively studied, but since initial findings in the early 1990s that TGF-β downregulates MHC I surface expression, this mechanism has been largely ignored. Due to the importance of MHC I expression in immune surveillance and intracellular pathogen clearance, identification of the TGF-β signaling pathways and target genes involved in antigen processing and presentation may facilitate the development of therapeutics to regulate MHC I expression during disease. In this study, we demonstrated that all three TGF-β isoforms are capable of downregulating constitutive MHC I surface expression and that this mechanism is conserved across species and cell types and dependent on Smad3 signaling. We then used RNA-seq in equine and human MSCs to identify possible target genes associated with antigen processing and presentation that were differentially expressed in TGF-β-treated MSCs. We found numerous genes associated with antigen processing and presentation were downregulated in TGF-β-treated MSCs with B2M and ERAP1 consistently downregulated in both species. These findings identified previously unknown gene targets of TGF-β and provide evidence that TGF-β downregulates MHC I surface expression through canonical Smad3 signaling.

The three TGF-β isoforms have very similar physical structures and all signal through the canonical Smad2/3 signalling pathway, but differences are known in the intrinsic activity of each isoform. For example, Yang and Kaartinen found that in vivo replacement of TGF-β3 with TGF-β1 was insufficient to completely correct the cleft palate phenotype of TGF-β3 knockout mice (Yang and Kaartinen, 2007). We are the first to report that like TGF-β1 and TGF-β2, TGF-β3 also downregulates MHC I surface expression. It is possible that the ability to downregulate MHC I expression by TGF-β is conserved among the isoforms due to its importance in regulating antigen presentation and inflammation; however, unlike TGF-β1, the importance of TGF-β2 and TGF-β3 in regulating MHC I expression in vivo has not been investigated. As MHC I expression increased after inhibition of the type I TGF-β receptor, ALK5, it is likely that endogenous TGF-β signaling plays a role in regulating constitutive MHC I surface expression. According to the Human Protein Atlas, RNA expression of TGFB2 is highest in the brain, eye, and reproductive organs, which are considered immune privileged tissues while TGFB3 also has high RNA expression in the brain and reproductive organs (Thul and Lindskog, 2018). The presence of TGF-β in sites of immune privilege likely contributes to the low MHC I expression in these tissues, which reduces autoantigen presentation (Niederkorn, 2003). The role of TGF-β2 and TGF-β3 in regulating inflammation and MHC I expression in vivo has not been explored in part due to the importance of these two isoforms in embryonic development. Homozygous knockout of TGF-β2 is a lethal embryonic mutation, so an inducible TGF-β2 knockout mouse model may be able to address how TGF-β2 regulates MHC I expression and immune privileged in vivo.

We used both equine and human MSCs to identify genes associated with antigen processing and presentation that were downregulated by TGF-β. Using RNA-seq and qPCR, we identified that TGF-β consistently downregulates B2M and ERAP1 in MSCs from both horses and humans. B2M was shown to be essential for MHC I surface expression after the first B2M knockout mice were discovered to be deficient in CD8^+^ T cells due to a lack of MHC I surface expression (Zijlstra et al., 1990). Downregulation of ERAP1 is also known to affect surface expression of MHC I (York et al., 2006; Wang et al., 2013). As ERAPI trims antigenic peptides to fit into MHC I binding grooves, decreased ERAP1 expression likely results in decreased peptide loading and empty MHC I molecules being too unstable to be expressed on the cell surface (Gruhler and Früh, 2000). It’s also well established that normal expression of TAP2 is necessary for MHC I surface expression (Ljunggren et al., 1990). While TAP2 was downregulated in some TGF-β-treated MSCs, it was not consistently downregulated in all samples or at all time points tested. TAP2 was significantly downregulated in all horses 12 hours after TGF-β treatment, but as TAP2 is critical for maintaining antigen processing and presentation, expression dropping below a certain threshold may activate TAP2 expression and override TGF-β signaling. MSCs treated will TGF-β always remain positive for MHC I surface expression and transcription factors like NLRC5 may play a role in activating transcription of genes targeted by TGF-β to maintain a minimum MHC I surface expression level (Meissner et al., 2010). While B2M is known to be positively regulated by NLRC5, TAP2 is not (Jongsma et al., 2019) so further research is needed to understand both the positive and negative regulatory networks that control antigen processing and presentation genes.

Many of the genes associated with antigen processing and presentation have similar promoter regions with binding sites for interferon response factors and NF-kb (van den Elsen et al., 2004; Compagnone et al., 2019) and ERAP1 is also known to be positively regulated by p53 (Wang et al., 2013). IRF1, IRF2, TP53, and genes encoding the NF-kb subunits were not differentially expressed in TGF-β-treated equine and human MSCs, however. Smad3 can directly inhibit gene expression by inducing chromatin remodeling through the recruitment of histone deacetylases, interfering with active transcription factor complexes, or by interacting with additional transcription regulators to inhibit transcription (Hill, 2016). Additional research at the chromatin level is needed to understand which of these mechanisms are utilized by Smad3 to inhibit expression of B2M and ERAP1. Genome wide approaches including ChIP-seq and ATAC-seq would be useful for identifying direct Smad3 targets or indirect targets that might be downregulated through a secondary or negative feedback mechanism.

We have established that TGF-β downregulates ERAP1 and B2M gene expression and that this likely contributes to downregulation of MHC I surface expression, but both B2M and ERAP1 have additional functions beyond simple antigen processing and presentation. ERAP1 is involved in suppression of inflammasome pathways (Blake et al., 2022) and cytokine receptor shedding (Cui et al., 2003) and polymorphisms in ERAP1 are associated with increased risk of ankylosing spondylitis (Roberts et al., 2017; Ma et al., 2022). As downregulation of ERAP1 is known to alter the pool of peptides presented via MHC I and subsequent immune responses (Hammer et al., 2007), further investigation into how TGF-β alters the MHC I-associated peptidome is warranted. Soluble B2M has been shown to trigger NLRP3 inflammasome activation in tumor associated macrophages (Hofbauer et al., 2021), act as an antimicrobial peptide in infection (Chiou et al., 2021), and promote neoplastic cell migration and metastasis (Chen et al., 2008; Wang et al., 2018). To our knowledge, no other studies have reported that ERAP1 is downregulated by TGF-β while Sun et al. is the only other study to demonstrate B2M is downregulated by TGF-β signaling. In this study, the authors showed that the addition of 10 ng/ml of exogenous TGF-β1 to cell cultures, reduced expression of B2M protein in ovarian cancer cell lines, which then decreased cell proliferation and migration (Sun et al., 2016). Further investigation of the role of TGF-β in regulating ERAP1 and B2M gene and protein expression may lead to strategies or therapeutics to target these genes in inflammatory diseases and cancer.

Using human and horse primary cells allowed for us to examine the variability between individuals, but using outbred species also makes it more difficult to interpret any changes in gene expression of the MHC I genes themselves. The α1 and α2 domains of the MHC I molecule are highly polymorphic, but there are also conserved regions in the α3 domain across MHC I genes where CD8 binding occurs (Takeshita et al., 1993). Therefore, unless the donor’s individual haplotype is known, qPCR primers will often amplify multiple MHC I genes and RNA-sequencing reads may also be incorrectly mapped to the genome (Brandt et al., 2015). Only one equine classical MHC I gene has been mapped to the genome, EQMHCB2, and while gene expression was downregulated in TGF-β-treated MSCs, the adjusted *p* value did not reach statistical significance. RNA from a larger sample size of horses was used in an attempt to confirm the RNA-seq for EQMHCB2, but the primers did not appear to be specific and produced multiple peaks in melt curves likely due to amplification of other MHC genes. All three HLA genes in TGF-β-treated human MSCs were downregulated, but only HLA-B had a Log_2_FC >1 and an adjusted *p* value < 0.05. Published qPCR primers for HLA-A, HLA-B, and HLA-C were tested, but similar to the horse samples, the primers appeared to amplify multiple targets. Inbred mouse strains where the haplotype and MHC I gene sequences are known could be used to determine if TGF-β downregulates MHC I genes, but the mouse MHC I genes are not necessarily analogous to human HLA genes. In addition to determining if Smad3 directly downregulates other antigen processing and presentation genes, ChIP-seq would be able to detect if Smad3 binds to any upstream regulatory regions for the HLA-A,B,C genes.

Through this study, we have identified that canonical Smad3 signaling is responsible for downregulation of MHC I surface expression by TGF-β and that B2M and ERAP1 genes are also downregulated by TGF-β. The precise molecular mechanism by which Smad3 signaling downregulates B2M and ERAP1 has not yet been established, but this pathway may be a potential therapeutic target for regulating MHC I surface expression in disease.

## 5 Ethics Statement

The animal study was reviewed and approved by the Institutional Animal Care and Use Committee of North Carolina State University.

## 6 Author Contributions

AB conceived, designed, performed the experiments, and prepared the manuscript. AB and LS analyzed and interpreted the data. AE also contributed to data acquisition and analysis. All authors read and approved the final manuscript.

## 8 Acknowledgments

We thank Drs. Jenny Ting and Yisong Wan for their feedback and Dr. Doug Antczak and Donald Miller for providing reagents and technical assistance. The authors would also like to thank the North Carolina State University CVM Laboratory animal resources staff for their help with animal care.

## 9 Data Availability Statement

The raw RNA-sequence data and normalized cells counts generated for this study can be found in the Gene Expression Omnibus (GSE207394 and GSE220294). All other data are available upon reasonable request.

